# Accuracy of programs for the determination of HLA alleles from NGS data

**DOI:** 10.1101/183038

**Authors:** Antti Larjo, Robert Eveleigh, Elina Kilpeläinen, Tony Kwan, Tomi Pastinen, Satu Koskela, Jukka Partanen

## Abstract

The human leukocyte antigen (HLA) genes code for proteins that play a central role in the function of the immune system by presenting peptide antigens to T cells. As HLA genes show extremely high genetic polymorphism, HLA typing on the allele level is demanding and is based on DNA sequencing. Determination of HLA alleles is warranted as many HLA alleles are major genetic factors that confer susceptibility to autoimmune diseases and is important for the matching of HLA alleles in transplantation. Here, we compared the accuracy of several published HLA-typing algorithms that are based on next generation sequencing (NGS) data. As genome screens are becoming increasingly routine in research, we wanted to test how well HLA alleles can be deduced from genome screens not designed for HLA typing. The accuracies were assessed using datasets consisting of NGS data produced using the ImmunoSEQ platform, including the full 4 Mbp HLA segment, from 94 stem cell transplantation patients and exome sequences from the 1000 Genomes collection. When used with the default settings none of the methods gave perfect results for all the genes and samples. However, we found that ensemble prediction of the results or modifications of the settings could be used to improve accuracy. Most of the algorithms did not perform very well for the exome-only data. The results indicate that the use of these algorithms for accurate HLA allele determination based on NGS data is not straightforward.

## Introduction

Successful transplantation of solid organs and stem cells requires good immunogenetic matching. The matching of alleles in HLA genes is particularly crucial (1,2). All HLA genes are in the MHC gene complex on chromosome 6p21.3 (3). The classical transplantation HLA genes, HLA A, B, C, DRB1-5, DQA1, DQB1 and DPB1, encode for integral membrane proteins, which bind short peptides that are recognized by T-cells of the cellular immune system. HLA molecules are the starting point of the adaptive immune response, making them interesting not only in the normal immune response but also in the susceptibility to autoimmune diseases and tissue and cell transplantation matching. The hallmark of the HLA genes is their very extensive genetic variation. More than 16,000 alleles have been identified in the HLA genes in the IPD-IMGT/HLA database (4). The determination of the exact allelic forms of HLA genes is warranted particularly in stem cell transplantation between unrelated persons (2). However, their determination is challenging due to the extremely high number of alleles, closely related gene sequences between HLA genes, and similar or even identical gene segments shared by some alleles or genes.

Currently, PCR-based molecular techniques, including Sanger sequencing, are the gold standard for HLA allele determination (“HLA typing”). Traditionally the regions of interest have been exons 2 and 3 for the class I genes HLA A, B and C and exon 2 for the class II genes DRB1-5, DQA1, DQB1 and DPB1. Methods based on next-generation sequencing (NGS) techniques are also emerging (5,6, 7). Most of the widely used high-throughput NGS sequencing instruments produce only short reads, from approximately 100 to a few hundred nucleotides per read. This, together with the lack of information on which of the chromosome pairs that a sequencing read originates from, causes problems in the bioinformatics analyses of NGS data. These problems are particularly substantial in the case of the highly polymorphic HLA genes, which can differ from each other only by a single nucleotide. Additionally, different genes may share highly homologous sequences, leading to problems in the short read alignment. NGS technologies producing longer read lengths have the potential to solve many of these problems.

NGS data may originate from different genomic sources, depending on the targeting method utilized. For example, a typical genomic target region may be the whole exome, selected exons or the whole genomic sequence of the target genes. In the context of HLA typing, only a subset of exons, or alternatively, the whole MHC region may be targeted (5–10). NGS can also be performed without targeting, as is done in whole-genome or transcriptome/RNA sequencing.

As algorithms or computer programs are a key factor in the utilization of NGS data, we selected a set of publicly available programs for the determination of HLA alleles based on NGS data and tested their accuracies using two datasets with different characteristics. One data set included the whole genomic MHC region as part of ImmunoSEQ sequencing platform (21) in 94 patient samples, while the other, a set of samples from the 1000 Genomes catalogue (11), targeted primarily exons. The best HLA-typing results for clinical transplantation use are likely to be achieved by using targeting panels specifically designed for HLA allele determination. It can be assumed that most alleles are accurately typed using these panels. However, genomic or exomic sequence data, including the MHC segment, are generated in research projects, that although they do not primarily aim at the determination of HLA alleles, would still benefit from valid interpretation of HLA alleles based on the NGS sequence produced. An example described in the present study is the use of the ImmunoSEQ sequencing platform (21), which primarily focuses on identification of rare variants in immunologically relevant regulatory areas and includes the full genomic sequencing of the MHC region. We selected both assembly-based and alignment-based methods, which increases methodological diversity. In addition, we used an ensemble approach (12) to test whether accuracy could be increased by combining results from different programs. The classifiers that are part of an ensemble should be diverse enough so that they do not all produce the same erroneous result. Hence, it was essential to select programs that applied different approaches, either assembly-based or alignment-based. We also describe some modifications to the default instructions to achieve more reliable results with one of the programs. While none of the programs, when used alone or according to the standard default instructions gave perfect allele assignments, our results show that the ensemble approach produced better results.

## Materials and Methods

### Finnish Red Cross Blood Service (FRCBS) dataset

A total of 94 samples from patients who were HLA typed for possible haematopoietic stem cell transplantation formed the FRCBS data set. This study was carried out in accordance with the recommendations of the Ethical Review Board of Helsinki University Hospital with written informed consent from all subjects. All subjects gave written informed consent in accordance with the Declaration of Helsinki. The protocol was approved by the Ethical Review Board of Helsinki University Hospital.

Clinical HLA-typing was performed from 2003-2008 in the HLA laboratory of the Finnish Red Cross Blood Service, using methods accredited by the European Federation for Immunogenetics. Four different techniques were used. For both low- and high-resolution HLA typing, LIPA (Innogenetics Group, Gent, Belgium), rSSO-Luminex technology (Labtype, One Lambda Inc. Canoga Park, CA, USA) and PCR-SSP (Micro SSP™ Generic HLA Class I/II DNA Typing _®_ Trays, One Lambda Inc. Canoga Park, CA; Olerup SSP® genotyping, Olerup SSP AB, Stockholm, Sweden) were used. The results were analysed with the appropriate software provided by the manufacturers. Sequence-based typing for determining the high-resolution HLA alleles was performed with AlleleSEQR PCR/Sequencing kits (Atria Genetics, Hayward, CA), using the ABI 3130xl genetic analyser (Applied Biosystems, Thermo Fisher Scientific, MA, USA), and the results were analysed with the Assign 3.5+ software (Conexio Genomics Pty Ltd, Fremantle, Australia) according to the supplier's instructions.

NGS sequencing was performed as part of the ImmunoSEQ V3 capture panel (21) using Roche SeqCap EZ Human MHC Design capture, which captures approximately 95% of the MHC/HLA region. The sequencing was performed with Illumina HiSeq 2000, yielding 100 bp paired-end reads and a median on-target coverage of 27.5x per sample. The data was quality checked using FastQC (FastQC) and adapters were trimmed using Cutadapt (13). For HLA genes, the coverages by ATHLATES were as follows: HLA A 14–138x; HLA B 13-99x; HLA C 14-165x; DRB1 22-216x; DQA1 14-191x; DQB1 15-165x; and DPB1 15-152x.

### 1000 genomes dataset

We used data of 63 samples from the 1000 genomes project (11), for which HLA typing had been carried out using Sanger sequencing (14). The samples represented three different ethnic groups: 33 Finnish, 15 Puerto Ricans, and 15 Yoruba. The HLA alleles for HLA A, B, C, DRB1 and DQB1 were available. Hence, these genes were tested. The samples were not selected based on the date of sequencing, depth of sequencing, or any other quality measure as our purpose was to test the methods on data produced using the standard, exome sequencing.

### Programs for determination of HLA alleles from NGS data

The programs can be categorized into read-mapping and assembly-based approaches. Instead of using a single reference, as is usually done for NGS data, the read-mapping methods are often based on the alignment of NGS reads with a reference sequence set consisting of all known HLA alleles. Then, the HLA alleles of the sample are predicted based on the properties of the alignments, often summarized as a likelihood or probability score formed by, for example, the number of mapped reads and the overall quality of the alignment. The assembly-based methods first construct or “assemble” the NGS reads into larger contigs, which are then queried against a reference database containing all known HLA allele sequences. The reference sequences and allele names are derived from the IPD-IMGT/HLA database (4).

Below, we briefly describe the methods selected for comparison in this study, but refer the reader to the references for further information on the programs.

All the methods were installed and used as instructed in their manuals.

#### ATHLATES

ATHLATES (15), version 1.0, is an assembly-based method developed for use with exome sequencing data. ATHLATES first filters the sequencing reads by aligning them to HLA allele sequences (either gDNA or cDNA) obtained from the IPD-IMGT/HLA database, allowing for soft-clipping to include intron-exon spanning reads. Reads mapping to more than one HLA gene (for e.g., HLA-A and non-HLA-A) are excluded. Paired-end reads are merged and reads with potential sequencing errors (low frequency k-mers) are discarded. Contig assembly is initiated by using each paired-end read as a contig. Contigs are then merged by considering those sharing an *l*-mer and *l* is decreased iteratively from the full read length until a fixed threshold is met. We kept track of the frequency at each base all the time. Contigs sharing longer and more high-frequency substrings are prioritized in comparisons and merging.

The exons of each HLA allele are then matched with the assembled contigs and the overall difference (Hamming distance) is calculated as the sum of the differences of each exon in the allele. Only alleles with no more than two mismatches and adequate coverage (20x read coverage, minimum of 85% of exon sequence covered by best-hit contigs, more than 70% of cDNA length captured by summed exon lengths) are considered further.

A list of candidate alleles is then formed by selecting those with no missed exons and no more than one mismatch. When the alleles correspond to multiple protein coding sequences, no mismatches are allowed. Pairs of alleles are formed from this list, scores are calculated for each pair (using a scoring scheme based on multiple sequence alignment), and, finally, the pair(s) with the best score is reported.

ATHLATES was installed and used as instructed in the ATHLATES User Manual 1.0. Novoalign V3.03.00 was used for alignment using the parameters recommended in the ATHLATES manual and for defining the fragment lengths according to the input data.

From the typing results, we selected only allele pairs listed under "Inferred Allelic Pairs”. As ATHLATES often reports more than one possible allele pair, the list was reduced to arrive at only one allele pair to make the comparison with other typing methods more unbiased. The selection was carried out by allowing the listed alleles to vote and then selecting the two lower-resolution alleles with the highest votes, e.g. A*11:01:01:01 and A*11:01:01:02 would result in allele A*11:01:01. Even if alleles at the lowest resolution level failed to achieve a higher vote than the others, the call was considered to be empty. If the files had no calls at all, they were interpreted as having a missing allele.

ATHLATES does not report only a single allele pair, but a list of the best allele pairs. To keep the comparison fair to other programs, a voting scheme similar to the one described below was used to select the most likely alleles. Selection of the most likely allele pair could also be performed, for example, based on population frequencies of alleles, but this approach was not utilized here.

#### HLAssign

HLAssign was developed (mainly) for HLA typing using specific targeted capturing with baits that was designed to consider the highly polymorphic sequences in the HLA region (15). The Windows 64-bit version 14.04 with database IPD-IMGT/HLA v 3.21.0 was used. HLAssign works by mapping all the reads to cDNA sequences from the IPD-IMGT/HLA database (4). It then discards those that are not completely covered or only have coverage on a small, central portion of the read. Several different parameters/statistics are then calculated for all the allele pairs formed from the remaining alleles. The allele pair with the highest weighted harmonic mean of the scaled parameters is selected as the most likely allele pair. This is the allele pair we used for comparisons.

#### OptiType

OptiType (16), version 1.3.1, is a mapping-based method and can produce typings from both DNA and RNA sequencing data. It also aims at utilising multi-mapping reads, whereas, in many other programs, these are discarded. OptiType gives results only for major class I genes (A, B and C). Even though in principle it should be possible to modify the program to also type class II genes, we did not carry out such modifications.

OptiType has its own HLA allele reference database, that is constructed from exons 2 and 3 of class I HLA alleles from the IPD-IMGT/HLA database (4). It also includes the intervening intronic sequences (using special phylogenetic-based imputation if intron sequences are missing) for use with DNA sequencing data. Non-classical HLA class I genes HLA G, H and J are also included. Sequencing reads are first aligned against the database, allowing multiple matches per read. A binary matrix is then created, indicating which alleles best align to each read. Using this matrix and integer linear programming, the best allele combination is selected by maximizing the number of reads mapping to each locus.

#### HLAreporter

HLAreporter version 1.03 (17) begins by filtering the NGS reads, which is done by mapping them to a reference sequence panel. This reference set consists of exons 2-4 for class I genes and exons 2 and 3 for class II genes. The sequences are obtained from the IPD-IMGT/HLA database (4). The panel is further modified by appending 50 bp of intronic sequence to both ends of each exon, including non-classical HLA class I genes. NGS reads that receive no mapping results or map perfectly to more than one gene are filtered away. The remaining set of reads for each gene is then used to assemble contigs. Then, to match the assembled contigs, two more databases are used: one with only exons 2 and 3 for class I and exon 2 for class II genes, and the other one with sequences for less polymorphic exons (exon 4 for class I and exon 3 for class II genes). Matching to the first database allows identification of the alleles, but often only at the G-group level, while the second database is used to further break down the result. Contig-HLA allele matching is performed by calculating scores for each contig (as a product of contig size, average depth of coverage, and percentage of exonic sequence) as well as counting the score for an allele by summing the scores of the contigs supporting this allele. The program only accepts perfect matches between the assembled contigs and the candidate HLA alleles. The best-scoring alleles are reported as the result.

A list of possible alleles may be produced, but there can be several possible alleles listed for both allele pairs. In the same way as for ATHLATES we used voting to reduce the lists to a single allele pair. If the voting gave no consensus, the gene call was marked as missing.

#### Omixon Target

This program, version 1.93, was the only commercially available program. Only a limited amount of information appears to be public and can be found on the www site of the producer www.omixon.com, where a reference to (18) is given. The algorithm (18) works by aligning reads against the IPD-IMGT/HLA sequences with certain constraints. It then scores the alleles and reports the most likely allele pair. However, the exact functioning of the program might differ from that described in the publication. As the Omixon Target program was no longer available by the end of year 2016 we also tested its updated version Omixon Explore (Version 1.0).

### Ensemble prediction

We used majority voting to combine the results from different typing programs into an ensemble prediction. Because of the multi-level nature of HLA alleles, there are situations where a gene is voted as two alleles, such as 01:01:01 and 01:01:02. In this case, the resulting majority vote would be 01:01, i.e., the most detailed level up to which the majority of the voting methods agree. To find such majority voted alleles, we used a tree to capture all the alleles and their votes. Let this majority voting tree be called V. The root node of V is an empty node, meaning no typing result (which can occur if there is no consensus on any of the alleles).

For each voted allele, a temporary voting tree is formed so that the level of detail in typing increases towards the child nodes. For example, allele 01:01:02 would result in a tree 01→01→02. Each node also tracks the given vote (by default one). Such temporary trees are then added under the root node of V so that new nodes are appended and, for the existing nodes, the vote count is incremented. Each node of V also keeps track of the alleles that have contributed to its votes.

For each sample and gene, all typing methods/programs are given two votes each, since the programs (typically) return an allele pair. However, for some loci there are typing results from only a subset of the programs (e.g., OptiType only types HLA -A, -B and -C). Therefore, there is not always the same number of voting programs for each gene. If a typing program returns an ambiguous result (for both or either allele), such results are skipped and not used in voting. Some programs might also return missing calls meaning that the missing typing results are considered as “evidence” for the allele being absent. Thus, they contribute a vote for a missing allele, even though such results might also be indicative of various typing problems (related to sequencing or algorithms).

Selection of the majority-voted alleles is done by traversing V from the root node towards child nodes, if there is at least one child node whose vote reaches the required threshold (i.e., more than half of all the votes). If more than one child node has the same (highest) number of votes, the child can be selected randomly. When it is not possible to go any deeper (due to going below the voting threshold or being at a leaf node), the allele represented by this node is selected, and the votes of all alleles that contributed to this node are subtracted from V. This selection is done as long as there are nodes that have received more than half of all the possible votes. If there are no nodes with sufficient votes to start with, the result is deemed ambiguous.

Depending on the threshold, the voting result can range from no alleles to several alleles. Varying the threshold can be used to control the number of returned alleles. However, it is not always possible to get, for example, just a pair of alleles as the result. This is due to possible ties in voting. In this case, either one or three alleles may be given as the result for any gene. In our comparisons, we used a threshold of 0.5. In the event of more than two alleles per gene, the genes were investigated to see if there were alleles that could be combined at the level of the accuracy of the reference typing. As the voting does not necessarily return only one allele pair per gene, varying numbers of allele-comparisons may have to be performed. For example, if for a given sample the gene voting resulted in three alleles, one of which could not be matched to the reference typing, the unmatched allele is counted as an error and thus the result is three allele comparisons.

## G groups

Some programs, such as the HLAreporter, do not return allele pairs, but instead report G groups. A G group for a gene consists of the alleles with identical nucleotide sequences in exons 2 and 3 (for HLA class I genes) or exon 2 (for HLA class II genes). A single G group can contain tens of different alternative alleles, reflecting uncertainty in the typing. For such results, a temporary G group tree is formed consisting of all the alleles in the G group and each node (also parent nodes) have weights of one. For example, for G group A*02:11:01G both alleles 02:11:01 and 02:69 would get one vote. The lower-resolution type 02 gets only one vote as well even though it is shared by both alleles in the G group. The G group tree is then added to the actual voting tree V.

When selecting a majority voted allele, the whole G group tree is again subtracted from the voting tree if the majority voted allele was voted by this G group.

## Results

We tested four publicly available programs (HLAssign, HLAreporter, ATHLATES and Optitype) and one commercial (Omixon) program package for their accuracy in the interpretation of HLA alleles based on genomic NGS sequence data. Two data sets were tested. First, the FRCBS data set (N= 94 samples of apparent Finnish origin) was produced using the ImmunoSEQ V3 platform (21) that has been developed to screen rare DNA variants in immunologically relevant gene regions and has not been optimized for HLA typing. The ImmunoSEQ platform uses the Roche SeqCap EZ Human MHC Design capture and basically produces the full genomic sequence of the 4 Mbp HLA segment, with nearly 100% coverage of exons of the HLA genes. Sequencing depths of HLA genes, as calculated by ATHLATES, were as follows: HLA A 14–138x; HLA B 13-99x; HLA C 14-165x; DRB1 22-216x; DQA1 14-191x; DQB1 15-165x; and DPB1 15-152x. The FRCBS data set had been previously typed using standard EFI-accredited clinical methods. Second, to test the exome-only data, we selected 63 samples from the 1000 Genome dataset; the samples were of Finnish (N= 33 samples), Puerto Rican (N= 15 samples) and Yoruba (N= 15 samples) origins. The 1000 genome data set covered only the exomes of HLA genes, rather than the entire genomic sequence of MHC.

### FRCBS data

The sequencing datasets were first trimmed for adapter sequences and nucleotides of poor quality using Cutadapt (13). All typing programs were run as described in the Materials and Methods. As one of the 94 samples obviously showed a loss of heterozygosity (see below for details for this sample), the comparisons were done with only 93 samples.

Not all the programs were intended to be used for the typing of all HLA genes. Hence, the comparisons were divided into HLA class I and II genes. Since they were designed for this purpose, all the programs were able to give typing results for HLA A, B and C genes. We calculated the concordance between the results from each program and the reference, clinical HLA result (Figure 1, Table 1) for each of the class I genes separately, as well as for class I combined. Overall, the best performing programs were ATHLATES, Omixon and OptiType, which all gave the same alleles as the reference typing in 99-100% of cases. The other two programs gave concordant results in 96-97% of the cases. The ensemble prediction was beneficial as 100% concordance with the reference typing was achieved with this method for all the three HLA class I genes

**Figure 1.**
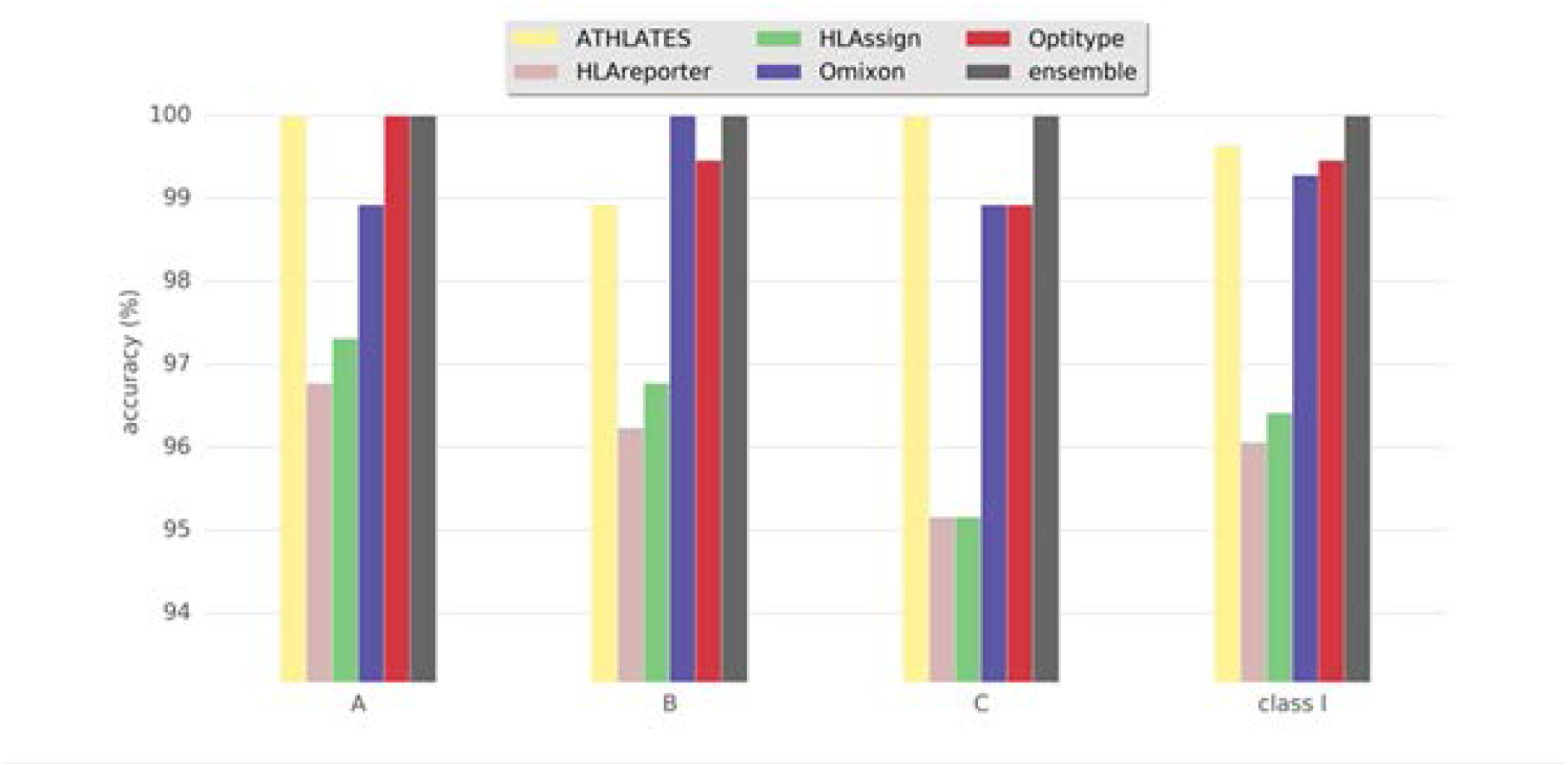
Accuracy of HLA interpretation programs to determine HLA class I alleles in the FRCBS data set comprising 93 Finnish individuals. Concordance rate to standard clinical HLA typing is shown for each program package and ensemble result.

**Table 1.** Summary of accuracies of the HLA interpretation programs and the ensemble result based on FRCBS data set comprising 93 samples whose full HLA segment was NGS sequenced using ImmunoSeq v3. Concordance rates to standard clinical HLA typing are shown.

|  | Program | Concordant result | Different | Accuracy % |
| --- | --- | --- | --- | --- |
| Class I | HLAssign | 538 | 20 | 96.42 |
|  | HLAreporter | 533 | 25 | 95.52 |
|  | ATHLATES | 556 | 2 | 99.64 |
|  | Optitype | 555 | 3 | 99.46 |
|  | Omixon | 554 | 3 | 99.28 |
|  | Ensemble | 558 | 0 | 100 |
| Class II | HLAssign | 703 | 45 | 93.98 |
|  | HLAreporter | 836 | 15 | 98.24 |
|  | ATHLATES | 703 | 147 | 82.24 |
|  | Optitype | na | na | na |
|  | Omixon | 822 | 28 | 96.71 |
|  | Ensemble | 848 | 4 | 99.53 |

Figure 2 and Table 1 show the results for class II genes. It is noteworthy that OptiType did not return any results for this set of genes; therefore, OptiType was not included in the final comparison. Even though HLAreporter was not among the best performing programs for class I genes, it performed best for class II genes, with a 98.24% concordance with the reference typing. Omixon also achieved an excellent result with a concordance rate of over 96%. It is noteworthy that ATHLATES gave a poor performance for the DPB1 gene. The program is clearly not designed for DPB1 typing using the default settings and instructions. However, as shown later, its performance can be enhanced significantly with certain modifications. DQB1 seemed to present a challenge for HLAssign, with a concordance rate of only approximately 80%. We did not investigate this further. The ensemble result for class II gave an excellent outcome with a 99.53% concordance rate. Four DRB3-5 alleles were not concordant. A DRB5*02:06; 02:01 heterozygosity was suggested in a sample with only one DRB5-associated class II haplotype and an extra DRB4*03:01N was proposed in three samples with DRB1*01;15 haplotypes. It should be noted that a discordant result is not be interpreted as incorrect, as it only implies discordance to the reference result.

**Figure 2.**
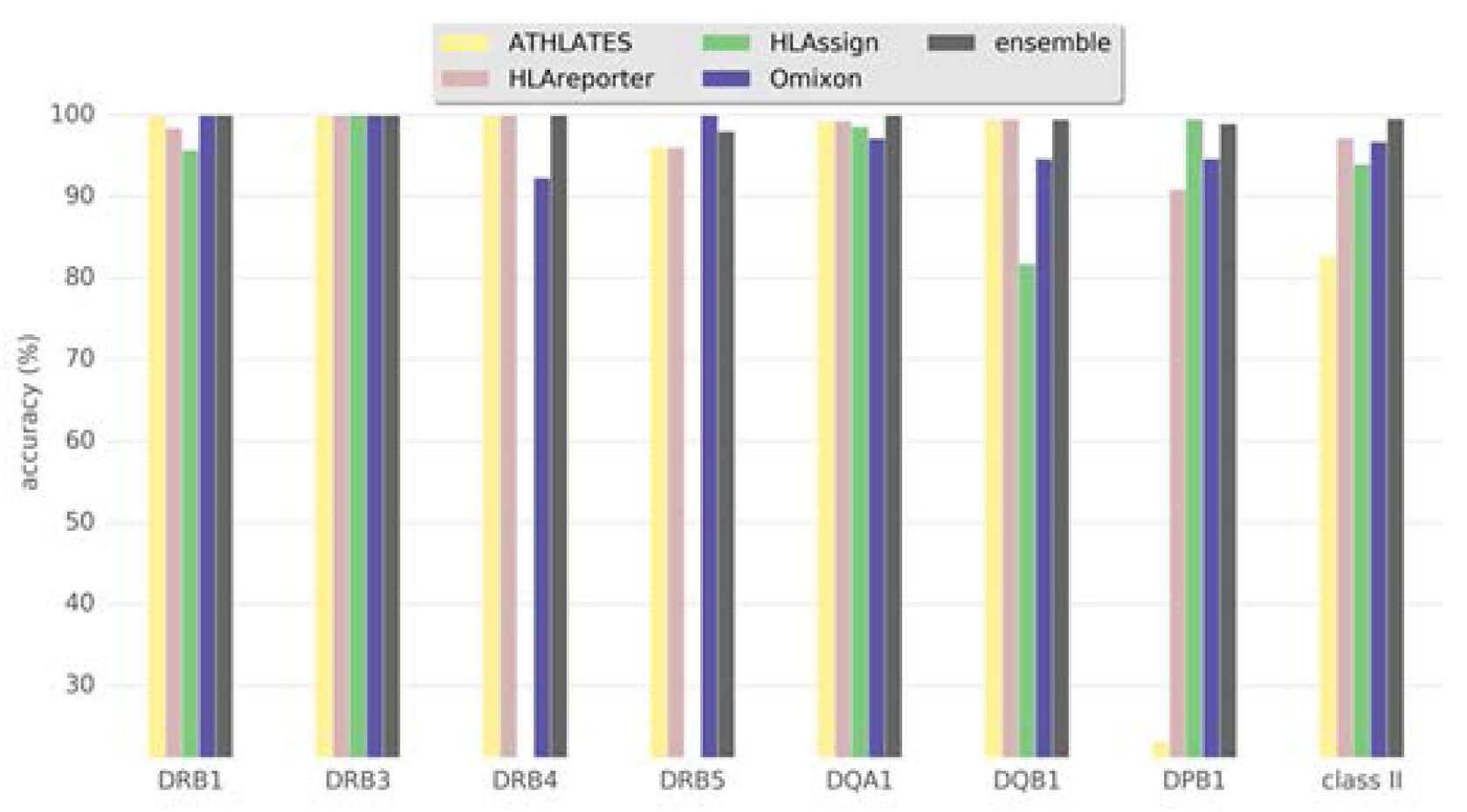
Accuracy of HLA interpretation programs to type HLA class II alleles in the FRCBS data set comprising 93 Finnish individuals. Concordance rate to standard clinical HLA typing is shown for each program package and ensemble result.

Since not all clinical HLA typings were performed to the level of a single allele with a unique amino acid sequence, we next tested whether the concordance rates would change if only those cases with 4-digit allele assignments were included in the analyses. For class I the accuracies were in general slightly lower, except for ATHLATES, which improved its accuracy. The differences seen in class II gene results were negligible (data not shown).

The discordant results varied between the different software. However, some samples seemed to be problematic in particular loci as the majority vote gave more than two possible assignments or the consensus could be drawn only at the two-digit level. As an example, there were three samples with a deviant majority vote result and a discordant HLA assignment with Omixon Target (Table 2). The discordant assignment could not be explained by low coverage or sequencing depth, as both alleles had an average coverage range of 97-100 and average sequencing depth range of 118-322. Furthermore, two (T5450 and T5684) out of three discrepancies disappeared after analysing the data with an updated version of the Omixon software, Omixon Explore. In the only remaining discrepant sample (T5703), the average sequencing depth of HLA DQB1*03:22 was lower than that of HLA DQB1*03:01 (118 versus 173, respectively). The full sequence of DQB1*03:22 allele is not known.

**Table 2.** Sequencing depth and coverage in three samples with discrepant results in HLA interpretation. Two samples T5450 and T5703 had an HLA DQB1 discrepancy and sample T5684 had an HLA-A discrepancy.

|  | Exons covered | Depth range | Exon 1 | Exon 2 | Exon 3 | Exon 4 | Exon 5 | Exon 6 | Exon 7 | Omixon Target | Omixon Explore | Reference method |
| --- | --- | --- | --- | --- | --- | --- | --- | --- | --- | --- | --- | --- |
| **DQB1 all samples** | 5 | 3-279 | 37-305 | 3-105 | 51-446 | 40-307 | 155-413 | na | na |  |  |  |
| **T5450** |  |  |  |  |  |  |  |  |  |  |  |  |
| **DQB1 allele 1** | 4 | average 134 | 167 | 89 | 148 | 178 | - | na | na | 02:01 | 02:01 | 02:01 |
| **DQB1 allele 2** | 4 | average 138 | 167 | 89 | 148 | 178 | - | na | na | 02:06 | 02:02 | 02:02 |
| **T5703** |  |  |  |  |  |  |  |  |  |  |  |  |
| **DQB1 allele 1** | 4 | average 173 | 189 | 77 | 245 | 205 | - | na | na | 03:01 | 03:01 | 03:01 |
| **DQB1 allele 2** | 2 | average 118 | - | 77 | 158 | - | - | na | na | 03:22 | 03:22 | 03:01 |
| **HLA-A all samples** | 7 | 16-322 | 4-206 | 5-239 | 4-305 | 4-463 | 12-426 | 1-389 | 1-366 |  |  |  |
| **T5684** |  |  |  |  |  |  |  |  |  |  |  |  |
| **HLA-A allele 1** | 6 | average 322 | 206 | 239 | 305 | 463 | 426 | - | 1 | 02:01 | 02:01 | 02:01 |
| **HLA-A allele 2** | 6 | average 322 | 206 | 239 | 305 | 463 | 426 | - | 1 | 02:197 | 02:01 | 02:01 |

Among the 94 Finnish samples, one was found to have a loss of heterozygosity (LOH) in the original clinical typings. Hence, it was not included in the comparison. In the clinical typings, three separate samples from the individual were required to confirm the heterozygosity in the HLA C locus. Obviously, higher proportions of malignant cells with an LOH hampered HLA determination. The NGS programs suggested homozygous HLA C assignment, which was also suggested by the majority vote. It is noteworthy that Omixon correctly suggested an unbalanced result in HLA C locus, indicating a minor fraction of reads from another putative allele. We have recently reported (19) a systematic screening of LOH in patients waiting for stem cell transplantation due to hematological malignancies. The sample from this study was not included in our previous study (19).

### Modified ATHLATES

Improvements in the ATHLATES results could be achieved by using the Mosaik v2.2.3 (20) aligner as described in the ATHLATES user manual 1.0 with the addition of the –om option to redirect multiple mapped reads to a separate bam file. When these reads were excluded in the HLA typing phase substantially better results were obtained (details not shown), demonstrating that minor modifications to the programs may result in more accurate performance.

### 1000 Genomes data

As exome sequencing is probably the most widely used method for genomic sequencing, we tested the ability of the programs to determine HLA alleles from such data. We used 63 samples from the 1000 Genomes Project for which HLA alleles had been determined using Sanger sequencing (14). The selected samples were from three different ethnic groups (33 Finnish, 15 Puerto Ricans, 15 Yoruba) and for each sample the HLA types for five genes (A, B, C, DQB1, DRB1) were available. Samples were not selected based on the date of sequencing, depth of sequencing, or any other quality measure as our purpose was to test the methods on standard, exome sequenced samples.

HLAssign was tested on several exome datasets, but it failed to produce calls, except for just a few genes in some samples. Thus, HLAssign was excluded from further analyses with the 1000 Genomes Project data. Additionally, we did not test the commercial Omixon program because we ran out of free trial credits. Its performance with 1000 Genome samples has been published (18).

For both HLAreporter and ATHLATES, we used voting to get a single best allele pair for each gene. If this did not produce a consensus, the allele call was marked as ‘no call’ and was counted as a typing error. If voting gave a low-resolution assignment, it was also marked as ‘no call’. Missing calls were also counted as errors unless the reference typings indicated a missing allele as well, although there were no missing calls for any of the typed genes in the 1000 Genomes set.

Results for the comparisons are shown in Table 3. To estimate the relative accuracy between the programs for the cases where data quality and/or coverage was not a limiting factor, we identified the intersection of genes for which the different programs returned results and counted the accuracies for the resulting set of alleles. The results are also shown in Table 3. Both HLAreporter and ATHLATES missed a large fraction of the alleles, whereas OptiType gave calls for all the alleles, with an accuracy of over 98%. This difference may be related to the fact that HLAreporter requires higher read coverage than was available. Additionally, the assembly-based methods cannot cope with sequence gaps, whereas gaps present no serious problem for the mapping-based OptiType. One factor explaining the HLAreporter discrepancy between the accuracies presented here and those achieved by Huang et al. (17) may be that we did not select only the good quality samples from the 1000 Genomes collection.

**Table 3.** Summary of accuracies of the HLA interpretation programs for 1000 Genomes samples. Missing allele refers to either a completely missing call or one for which no consensus could be formed based on a list of detected alleles. Intersection denotes the set of alleles for all sample-gene pairs that had a non-missing typing in all tested typing programs. No results could be achieved using HLAssign.

| Program | Number of alleles | Correct (all alleles) | Accuracy (all alleles) | Missing alleles (%) | Correct (intersection N=164) | Accuracy (intersection) |
| --- | --- | --- | --- | --- | --- | --- |
| HLAreporter | 630 | 202 | 32.06 % | 190 (30.16%) | 91 | 55.49 % |
| ATHLATES | 630 | 468 | 74.29 % | 78 (12.38%) | 137 | 83.54 % |
| Optitype | 378 | 372 | 98.41 % | 0 (0.00%) | 161 | 98.17 % |

## Discussion

A major finding in the present paper was that none of the programs alone gave perfect assignments for all HLA genes based on sequencing data that was not specifically designed for HLA typing. In fact, some of the programs were clearly unsuitable for use with certain genes, particularly when tested using the 1000 Genome data containing only exomes. Second, none of the programs consistently outperformed the other programs in all the cases and genes. Therefore, it was not possible to conclude the best performing program. Finally, with the ensemble prediction method, the accuracy improved, reaching almost 100% concordance with the reference. However, it is noteworthy that the ensemble approach is not very practical in clinical settings, but it may be used for research purposes to get the best possible HLA determination. In cases with discrepancy in typing results we could not conclude which of the allelic alternatives was the correct one, as it was not possible for us to have genuine control alleles by performing clinical grade NGS sequencing of the samples. Reliable HLA results can hence be achieved by using several programs and by applying the ensemble approach. The ensemble approach, however, has its drawbacks. All methods returning the same allele may lull us into a false sense of security regarding the allele call. It should be noted that even a unanimous result is not necessarily the right one, but can, for example, be caused by problems in sequencing or in the reference allele database. Use of sequencing methods that are based on short sequencing reads results in sequence alignment problems when applied to allele interpretation of the highly homologous HLA alleles and genes. We also noted that some of the problems with accuracy were associated with the properties of the software. A good example is shown in Table 2 in which the updated version of the Omixon program could resolve two of the three discrepancies found using the older version. Furthermore, this example shows the problems related to partial sequences known of some alleles: only exons 2 and 3 sequences of DQB1*03:22 are known. We were also able to show that the ATHLATES program could be adjusted to give much more reliable results by using modifiers as described by Lee et al. (20). We assume that similar modifications could be done to other programs as well.

The results presented here for some of the programs were likely worse than the results that might be achieved by an experienced HLA professional looking at the list of reported alleles and making interpretations based on the population allele or haplotype frequencies. We used and compared the programs as instructed by their manuals and applied default parameters. In addition, in cases where a program gave multiple alternatives with equal scores, we did not utilize the reference typing or population frequencies to determine the apparently correct alternative because there is usually no reference results available and the programs are used to determine the alleles. It is also possible that adjusting some settings may render the programs better suited to the study population, which might yield different, perhaps better, results.

Some differences in the relative performances of the programs might be due to the various versions of the IPD-IMGT/HLA database they use. However, we did not try to unify the versions since the programs use their own, slightly modified format of the database and because the programs were all released within a short period. The IPD-IMGT/HLA database is an obvious choice for reference. However, the sequences therein have been found to contain some errors (8) and we do not know whether some of the programs utilize the corrected sequences.

Determination of HLA class I alleles from the 1000 Genomes exome NGS sequencing data has been reported earlier (18). They reported an accuracy of over 90% between the NGS and Sanger data. In the case of the 1000 Genomes samples, it is notable that in our study most of the tested programs returned a significant number of missing results. A missing result may indicate that the gene was missing or that the quality of the data was insufficient for the program. The latter is a known problem because the targeting baits for exome sequencing are usually designed for the human reference genome, ignoring the very high variability in the HLA region (8). However, OptiType retained a very high accuracy even for exome-only data. Therefore, exons 2 and 3 for class I genes were likely well captured in the exome sequences of the 1000 Genomes Project. Alternatively, even though the exons might not be fully captured, OptiType is still able to make the most of the data, whereas some other programs had filtering criteria that could not be met when the exons were sequenced only partly. One such program is HLAssign where the cDNA allele sequences need to be fully covered in the alignment.

The results of the present study clearly indicate that selecting a program for HLA allele determination based on NGS data that was not designed for HLA typing purposes is not simple This process requires a good understanding of the type of NGS data produced and the HLA frequencies in the study population. Some modifications in the programs, the adoption of an ensemble approach or testing with multiple programs may be needed for accurate performance.

## Acknowledgements

This study was funded by Tekes, the Finnish Funding Agency for Innovation, as part of the SalWe GetItDone program and by Finnish government VTR funding for medical research.

The authors want to note that Omixon Ltd gave their commercial Omixon Target program free of charge for this study. Omixon has not participated in the interpretation of the results and has not seen them before publication.

